# Complex Traces: Using a Complex Adaptive Systems approach to examine morbidity and mortality among 19th Century migrants to South Australia

**DOI:** 10.1101/2025.02.19.639022

**Authors:** Angela Gurr, Matthew Brook O’Donnell, Alan Henry Brook

## Abstract

In multidisciplinary research interpreting interactions between diverse data sources requires a Complexity approach. A Complex Adaptive Systems (CAS) framework allows the relationships of multiple factors to be explored and may provide a more holistic and nuanced understanding. This study is innovative in explaining the potential benefits in a CAS approach to combining bioarchaeological and historical data when examining a rare archaeological skeletal sample of early migrants to South Australia (SA).

Macroscopic, radiographic and micro-CT methods were used for the analysis of the skeletal remains of a group of 19^th^ century migrants buried in an unmarked area of St Mary’s Anglican Church Cemetery. The relevant historical records explored were from British emigrant ships to SA (1836 to 1885) and the Church burial records (1847-1885).

Evidence of poor oral and general health was present in the skeletal material. Dental developmental defects indicated earlier health insults. Pathological manifestations in bone were compatible with joint and infectious diseases, and metabolic deficiencies. Historical documents recorded that the voyages to SA were challenging, with some ships experiencing a high death rate. Diseases e.g., measles and scarlet fever, and diarrhoea were frequently recorded as causes of death at sea for both subadults and adults. In the Colony, burial records showed similar causes of death for subadults, but for adults, accidents and tuberculosis were often reported.

The CAS approach provided insights beyond those from analysis of the individual sources. It increased understanding of emergent, non-predicted outcomes that resulted from interactions between multiple factors e.g., the impact of fluctuating economy, political instability and ideological pressures, on the health of migrants. The CAS framework is a valuable methodology for interpreting health patterns and can be further developed for a range of historical and contemporary health contexts.

## Introduction

### A Complex Adaptive Systems Approach

Multidisciplinary research that integrates and interprets different types of data from diverse sources can be challenging. These challenges are seen across a range of scientific and other disciplines. Different methods have been developed to address them [1], such as the Complex Adaptive Systems (CAS) framework approach, which has become more widely adopted [2–8]. The application of this approach, particularly within medical domains, to analyse findings from varied sources and investigate their relationships may provide a more holistic and nuanced understanding [2, 4, 7].

The characteristics of a Complex Adaptive System (CAS) are seen with the formation, development and ongoing functions of a social group, such as a new migrant colony [7]. Key components of a social CAS framework include interactions at multiple levels e.g., between individuals, small groups, families, neighbours, companies and government structures in networks at multiple timescales that give rise to emergent properties i.e., outcomes that cannot be predicted or anticipated from the interacting parts. The value of this perspective is that it can be used to guide the selection, organisation, analysis and interpretation of various data sources and may provide a fuller context for these findings, enhancing understanding of individuals and groups studied.

A vital tool that guides research thorough progressive stages or steps is the Wisdom Hierarchy [9]. First raw data collected from different sources are grouped or categorised. This produces information. Information is analysed to provide new knowledge, which is then interrogated and integrated to enhance understanding. It is in this final step of the Wisdom Hierarchy that key aspects of CAS are used to look at the interactions between the different sources of knowledge. The CAS approach results in new insights and subtleties beyond that arising from the independent treatment of each source. In Fig. 1, we develop further a Wisdom Hierarchy.

**Fig. 1.**
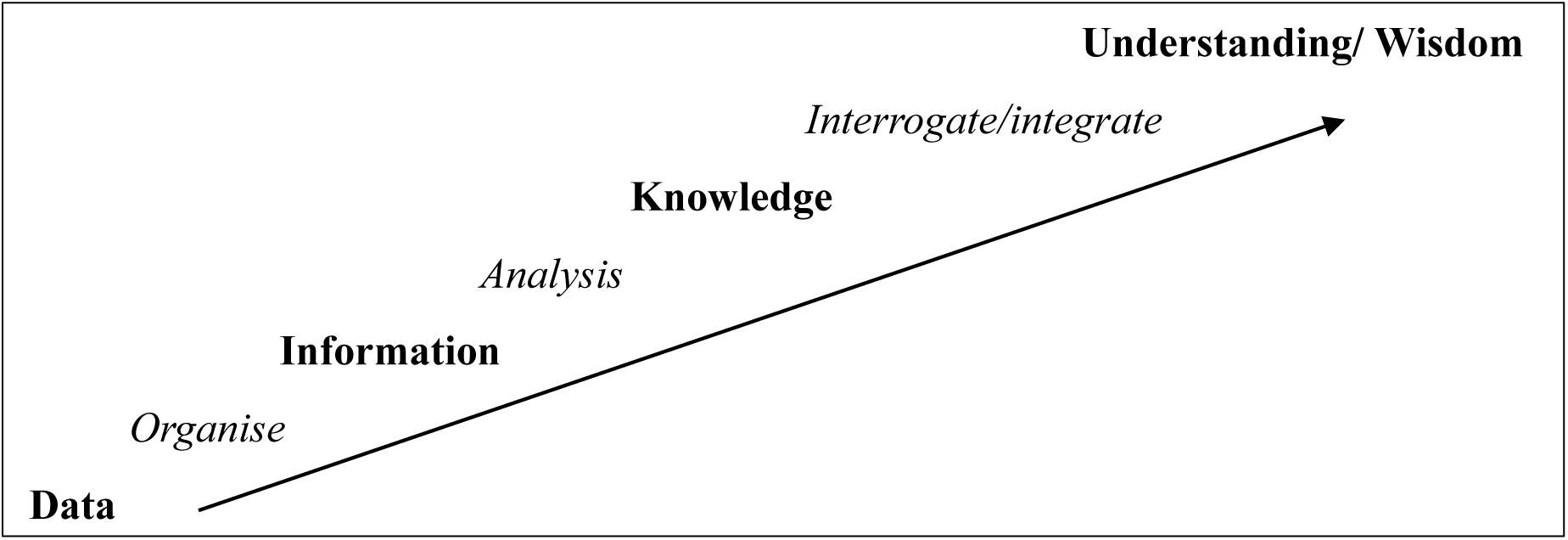
The Wisdom Hierarchy – a progression of steps from raw data is categorised and organised to create information. This is analysed to produce knowledge. Knowledge is interrogated to produce understanding. Further developed from Brook and Brook O’Donnell [10].

This study investigates multiple sources of data e.g., findings from the examination of archaeologically excavated human skeletal remains and diverse historical records, to understand the challenges faced by early migrants during the voyage and after arrival in the new colony of South Australia (SA). The years investigated are 1836 to 1885. These dates cover the initial period of *establishment* (1836-1848), followed by *settlement* of the province (1849 - 1866). The colony *matured* during the subsequent years (1867 - 1885).

#### Background for the new colony of South Australia

The American Revolution and independence from British rule (1773-1775/6) had caused political and commercial instability in Britain, arising from the loss of those colonies [11–13]. In Europe, the French Revolution of 1789-1794 brought civil unrest, social upheaval, and radical political changes [14, 15], while the war with Napoleon had required substantial capital and resources [16, 17]. The British aristocracy and political establishment were now greatly sensitized to the dangers of revolution by an oppressed populace. This was exemplified by the excessive use of force against a peaceful public protest that resulted in the Peterloo massacre (1819) [18].

The underpinning concepts and the formal establishment of the Provence of South Australia are well documented. Edward Gibbon Wakefield’s theoretical publication in 1829 proposed a systematic approach to establishing new colonies [19–21]. In these, rather than transporting convicts, voluntary emigration would be encouraged, and the British class system could be reproduced. Such new colonies could offer fresh opportunities for those affected by the gross overcrowding in the cities and towns of Britain and the greatly increased inequalities resulting from both international and national events.

The migration of workers from rural areas of Britain to urban industrial centres from approximately 1760 to 1850 followed the introduction of mass manufacturing of goods in factories using machinery [22–24]. Influxes of people in towns and cities without the infrastructure to cope produced overcrowding, unsanitary conditions, and the spread of disease [23–25]. Air and water pollution from factories adjacent to the workers’ housing also affected the health of the surrounding population [25–27].

During this period the coast and rivers of South Australia were being explored by Matthew Flinders (1801-1803) [28, 29], Charles Sturt (1828-1831) [30], and Collette Barker (1831) [31]. This region was seen as highly suitable for a new colony. With commercial backing in place from the formation of the South Australian Company [32, 33], the British Parliament passed the South Australian Colonisation Act (1834) [34]. An emigration fund, suggested by Wakefield (1829) [21], for the conveyance of healthy young migrants and their families to a new colony was planned and the ‘assisted passage’ scheme, sponsored by the Colonial Commissioners became available for migrants who were able to comply with the selection criteria [20, 34–38].

The first emigration ships for South Australia left Britain in 1836 [39–41]. These ships often carried different classes of passengers – e.g., cabin, intermediate and steerage/ assisted passage and had a Superintendent Surgeon to attend to the health of the migrants during the long voyages. The Passengers Act of 1835 [42], states that any ship conveying more than 100 people must have “some person duly authorised by law to practise in the UK as a Physician or Apothecary…taking with him a Medicine Chest and a proper supply of medicine, and instruments…”. This position was independent from the ship’s crew, the Surgeon being employed by the Colonial Commissioner. The logs kept by these Surgeons provide a valuable but under explored data source of the health of the migrants during the voyage.

Arrival in the new Province of South Australia did not end the challenges faced by the new migrant settlers. General environmental factors such as insufficient employment, a fluctuating economy and political instability [38], could have influenced their economic status and health as well as the eventual location of their burial.

While these historical events and decisions provide the necessary context for the migrant colony under investigation, viewing their interactions and the emergent outcomes they produced from a CAS perspective adds considerable value. There were short, medium and long-term outcomes from the interactions of the various decisions and actions, driven by ideological, economic, and political factors that were not, and likely could not have been, predicted at the outset. These outcomes could be either positive and negative and affect different individuals in different ways, the outcome in large part relating to individual resilience and the context. Therefore, the aim of this study is to explore if there are benefits in a CAS approach to combining bioarchaeological and historical data when analysing a rare skeletal sample of migrant settlers to SA.

## Materials and Methods

### Data sources

A summary of the different data sources used in this study are given in Table 1.

**Table 1:** A summary of the data sources investigated in this study.

| Data Source |  | Type of Data | Date Range |
| --- | --- | --- | --- |
| 1 | **Bioarchaeological**<br>St Mary's Cemetery Human Skeletal Material | Biological data from:<br>Macroscopic &<br>Radiographic examinations,<br>Micro-CT scanning | 1847-1927 |
| 2 | **Historical Documents**<br>Ships Records<br>Government<br>Ship's Surgeon's Logs: | Documentary | 1836-1885 |
| 3 | **Historical Documents**<br>St Mary's Church Burial Records | Documentary | 1847-1885 |

#### 1. Bioarchaeological investigation of human skeletal remains

##### Ethics

St Mary’s Anglican Church requested the excavation of the unmarked section of the cemetery and approved the study of this rare historical skeletal sample. Flinders University Social and Behavioural Research Ethics Committee provided ethics approval (SBREC Project number 8169). As this sample consisted of historical skeletal material of individuals whose identity are not known, there was no possibility or requirement to obtain informed consent.

##### Skeletal sample

The skeletal remains of 70 individuals (20 adults and 50 subadults - all migrant settlers to South Australia), were excavated from the unmarked section of St Mary’s Anglican Church Cemetery, near Adelaide, in 2000 [43]. The burials in this section of the cemetery took place between 1847-1927, but previous studies have shown that most of the burials in the unmarked section took place during the first decades after the establishment of the colony in 1836 [44–46].

The skeletal remains were accessed for research purposes from the 5^th^ of January 2021 to 31^st^ of November 2023, and between 22^nd^ January and 25^th^ July of 2024. Data derived from the macroscopic examination of the teeth and bones of the St Mary’s sample, and the radiographic examination and micro-CT scanning of just the dentitions were used [45–47].

Details of the scoring systems, and standards followed for estimation of age range, determination of sex, and the identification of pathological manifestation of disease are available from Anson [43] and Gurr et al., [45–47]. Criteria for the identification of, and systems followed to categorise tooth wear and/or pathologies related to oral health such as evidence of caries, and periodontal disease, and to measure dental developmental defects e.g. enamel hypoplastic defects and interglobular dentine, can be found in Gurr et al. [47].

Information and setting for the radiographic and Micro-CT scanning systems used e.g., Small Volume Micro-CT system - Bruker SkyScan 1276 [48], and the Nikon XT H 225 ST Large Volume Micro-CT system [49], as well as the different setting are published by Gurr et al., [47, 50].

#### 2. Historical documentation - Ships records

This study investigates two types of historical ships records, the government records and the ship’s Surgeons logs. Firstly, available documents relating to the emigrant ships that carried more than ten passengers from the United Kingdom (UK) to South Australia (SA) from 1836 to 1885 [51, 52], to provide information about the number of passengers and the conditions onboard e.g., the Summary Report and the Certificate of Final Departure [52, 53].

The documents completed by the Superintendent Surgeon assigned to each migrant ship from 1849 to 1865, as published in the South Australian Government Gazette [54], were used as they often recorded the cause of death for passengers who died at sea. The time periods covered by these two historical data sources differ slightly (Table 1) due to the availability of the data sources.

#### 3. Historical documentation – St Mary’s Church Records

Parish burial records for St Mary’s Anglican Church Cemetery, SA, began in 1847 [55]. To reflect the temporal period of the emigrant ships’ logs i.e., 1836-1885, the burial records have also been analysed until 1885. These documents provide insights into the morbidity and mortality during the early decades of the colony for a specific group of migrant settlers.

These burial documents record that N=143 individuals were interred *either* in i) the unmarked section of the cemetery often written as the ‘free ground’, ‘common ground’, or ‘unleased ground’, *or* ii) the location of their burial was not recorded, *or* iii) the location of their burial is unknown due to damage to the church records.

## Results

Table 2 provides an overview of the analysis and types of results obtained from the three data sources. The key variables from each source and how they were prepared for analysis and modelling are summarised.

**Table 2:** A summary of the different data sources used in this study, the key variables from each source and how they were prepared for analysis and modelling.

| Data Source |  | Unit of Analysis | Key Variables | Analysis & Modelling |
| --- | --- | --- | --- | --- |
| 1 | **Bioarchaeological**<br>St Mary’s Church Cemetery - human skeletal material | Individual | Age, sex, dental development defects, tooth wear, caries, periodontal disease, antemortem tooth loss. Skeletal pathologies, joint diseases & metabolic disturbances | Age group, descriptive observations, potential causes & future health implications |
| 2 | **Historical Documents**<br>Ships records<br>Government records<br>Ship’s Surgeon’s logs: | Voyage<br>Passenger | Date, count data by passenger type, number of deaths.<br>Year of death, Age, Cause of death. | Aggregation by time period & time series/trend analysis<br>Aggregation by cause of death & time series/trends analysis |
| 3 | **Historical Documents** | Individual | Estimated age, Cause of death | Grouped counts & aggregation by time period |
|  | St Mary's Church Burial records |  |  |  |

### 1. Bioarchaeological investigation of human skeletal remains

The demographic profile i.e., estimation of age range and determination of sex of the excavated skeletal remains from the unmarked section of St Mary’s Cemetery is available in the supplementary material as Table S1. It shows that many of these individuals were infants and subadults: This sample consisted of 21 infants of unknown age, 11 infants of 0-11 months, 19 subadults between 1-18 years and 19 adults of 19-50+ years of age. The identity of the 70 individuals is not known due to the lack of identifying markers such as gravestones. The multiple bioarchaeological techniques used for the analyses of the skeletons and dentitions (i.e., macroscopic, radiographic, and large and small volume micro-CT scanning), identified evidence of poor oral and general health conditions, as well as signs of earlier health insults in the form of dental developmental defects (Fig. 2).

**Fig. 2.**
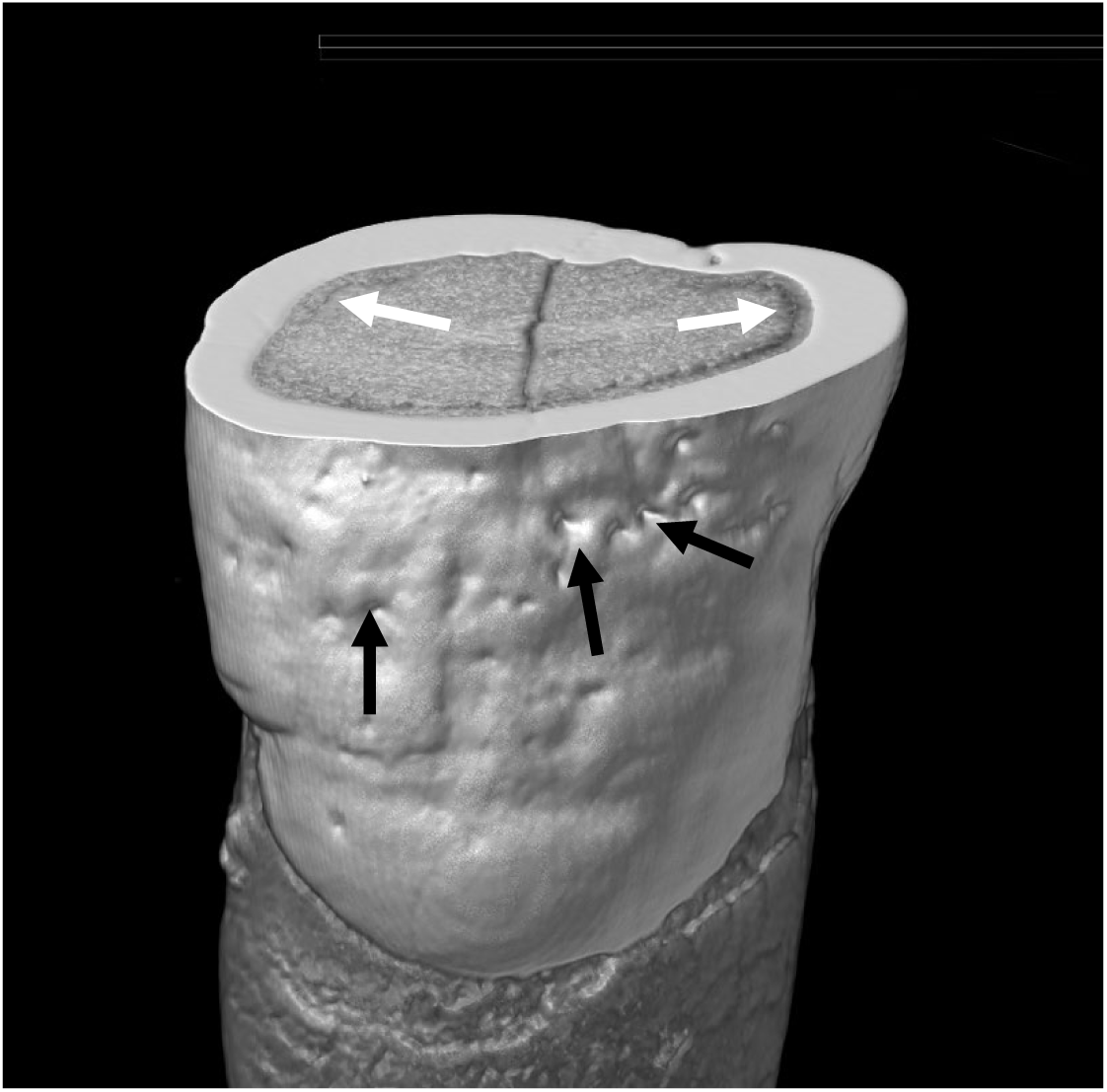
Small Volume Micro-CT – An image of a human tooth with developmental defects. This transverse (cross-sectional) view of the exterior and interior surfaces of a permanent upper incisor. Enamel hypoplastic defects (black arrows) and areas of interglobular dentine (white arrows) which are mineralisation defects of the dentine are highlighted.

Pathological manifestations and changes to the teeth and bones due to disease and/or deficiencies are listed in Table 3. Column 1 of Table 3 lists the health conditions that affected individuals in this group, with column 2 listing the observed evidence for the condition or defect. Column 3 provides information describing the potential outcome of such a condition/s on the health of the affected person.

**Table 3.** 1847-1927 – St Mary’s Cemetery skeletal sample - Evidence of previous health conditions and insults observed on teeth and bones.

| Condition/s affecting: *Adults* |  |  |
| --- | --- | --- |
| Condition/s | Observed Evidence | Potential causes and/ or implication for future health |
| **Poor Oral Health** | Caries –extensive | Poor oral hygiene - dentine/ pulp exposure – pain/infection, tooth loss |
|  | Dental abscess/es | Poor oral hygiene -chronic infection, pain, tooth loss |
|  | Calculus –build-up/ bone recession | Poor oral hygiene - gingivitis/periodontal disease/ systemic health issues |
|  | Periodontal disease | Tooth loss/ systemic health issues |
|  | Antemortem tooth loss | Inability to eat certain food types - reduced nutritional intake/ systemic health issues |
|  | Smoking – pipe notch | Change oral biome – increase systemic health issues |
| **Extreme Tooth Wear** | Diet/ health related- coarse food/ grinding teeth/GORD | Dentine exposure - pain- infection - tooth loss |
|  | Environmental e.g. abrasion/ attrition/ erosion | Loss of dental tissues – increase caries/ poor oral health, systemic health issues |
|  | Using teeth as tools | Trauma to teeth – infection – pain- tooth loss |
| **Joint Disease** | Vertebral osteophytes/ eburnation/ Schmorl’s nodes | Repetitive hard labour of a long period of time, reduced mobility– pain- disability |
|  | Morphological changes in bones - observed: radial notch of ulna, trochlear, olecranon, acetabulum, femoral head, talus head, navicular, & calcaneus | Reduced movement of the joints– pain- disability |
| **Dental Developmental Defects** | Enamel hypoplastic defects - | Reduction or cessation of tooth enamel production, evidence of systemic ill health during dental development |
|  | Interglobular dentine | Reduction in or cessation of the mineralisation of dentine, evidence of systemic ill health during dental development |
| **Condition/s affecting: *Subadults*** |  |  |
| **Metabolic Deficiencies** | Folic Acid – Vitamin B9 deficiency during pregnancy | Spina bifida occulta – failure of neural tube to completely close – mild – vulnerability in the posterior section of the sacrum |
|  | Vitamin. C deficiency – | Inflammation, bleeding, poor quality & quantity collagen – pain- potentially fatal - porous lesions in the cortical bone |
|  | Iron deficiency - | Fatigue- lack of oxygen to brain & muscles – weakness. Porous lesions in the cortical bone of the orbital roof |
| **Infectious Disease/s** | Periosteal reaction/ new bone growth | Chronic infection-inflammation- pain – poor systemic health – potentially fatal depending on condition |
|  | Congenital syphilis/ Dental morphological changes/ abnormalities |  |
|  | Tuberculosis/ Destructive lesions of the vertebral bodies & other bones |  |
| **Ingested Toxins** | Mercury - dental morphological changes/ abnormalities | Reduction or cessation of tooth enamel production, systemic ill health due to toxins. |
| **Dental Developmental Defects** | Enamel hypoplastic defects | Reduction or cessation of tooth enamel production, evidence of systemic conditions during dental development |
|  | Interglobular dentine | Reduction of the mineralisation of dentine, evidence of systemic conditions during dental development. |

Note: Further details of the conditions listed in Table 3 are available in previous studies [43, 45-47, 56, 57]

### 2. Ships records

#### Number of voyages and passengers over time -1836–1885

In total there were 885 voyages carrying more than ten passengers from the UK to SA from 1836 to 1885. The number of ships arriving in SA each year are shown in Fig. 3. The dashed line shows the accumulation of passengers from these voyages using a rolling 10-year window as a way of illustrating the impact of new arrivals in the growing colony over this time. There are no recorded emigrant ships arriving in SA for 1861, 1868 and 1872 and a single voyage in each of 1871 and 1885 [51, 52]. The busiest years were: 1849 (83 voyages), 1850 (76), 1854 (64), 1853 (60), and 1855 (57), all within the second period (1849-1870) (Fig. 3).

**Fig. 3.**
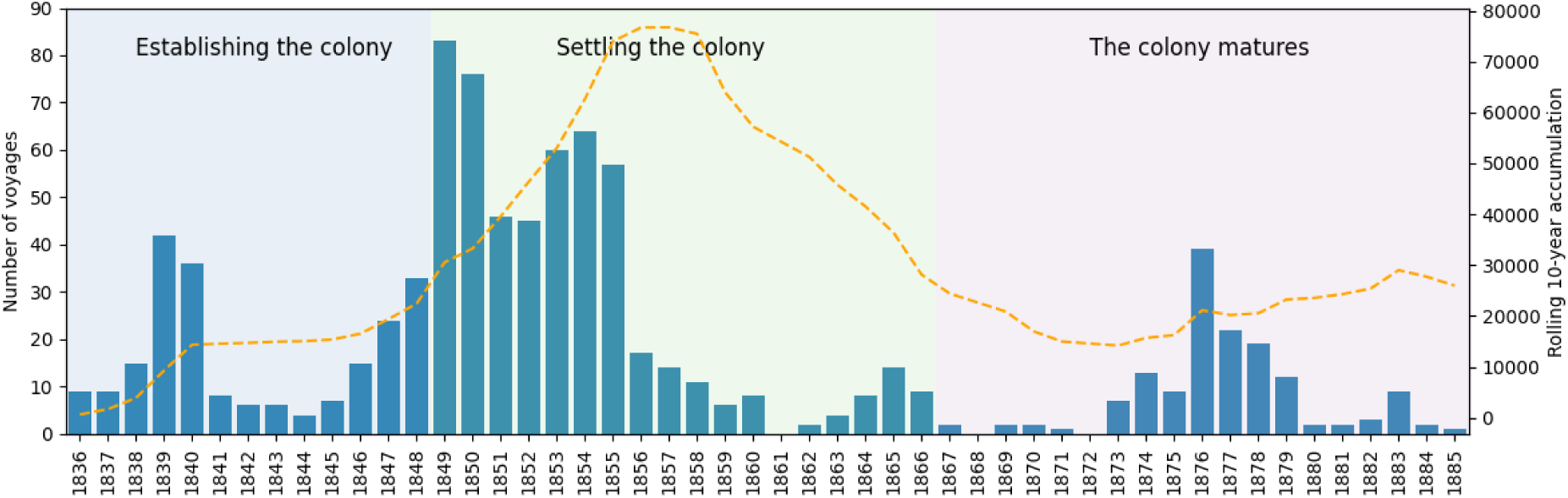
1836 - 1885. Number and distribution of emigrant voyages, carrying more than ten passengers from the United Kingdom to South Australia, in relation to the year of arrival. [51, 52]. Cumulative addition of individuals to the colony shown in the orange dashed line using a 10-year rolling window.

During the first period (1836-1848), in which the colony was being established, the average number of voyages per year was 16.5 (SD 13.0), driven by two key years 1839-1840 followed by a return to ten or less voyages per year until 1846. This is reflected in the flat accumulation line in Fig. 3. The second period (1849-1866) saw the settling and stabilization of the colony with a high demand for workers in the initial period up until 1855. The average number of voyages per year during this period was 29.1 (SD 28.2). However, dividing this period into an early section (1849-1855), with an average of 61.6 (SD 14.2) voyages per year and a later section (1856-1866), with an average of 8.5 (SD 5.3), better captures the complex dynamics of the settling period.

The accumulation of new arrivals (over a ten-year period) peaks towards the middle of the second period (around 1857) fitting with communities becoming established and organised. The final section (1867-1885) sees the colony maturing and requiring fewer external workers to sustain growth and development. This is reflected in the lowest average number of voyages per year of 7.7 (SD 10.0).

#### Number and type of mortality at sea over time - 1836–1885

There were many risks to the health of the 19^th^ century migrants bound for South Australia with some passengers dying during the long voyage. The number of infant, subadult and adult deaths per year from 1836 -1885 as recorded for British emigrant ships (with more than 10 passengers) to South Australia is shown in Fig. 4. The most vulnerable age groups were infants and younger children.

**Fig. 4.**
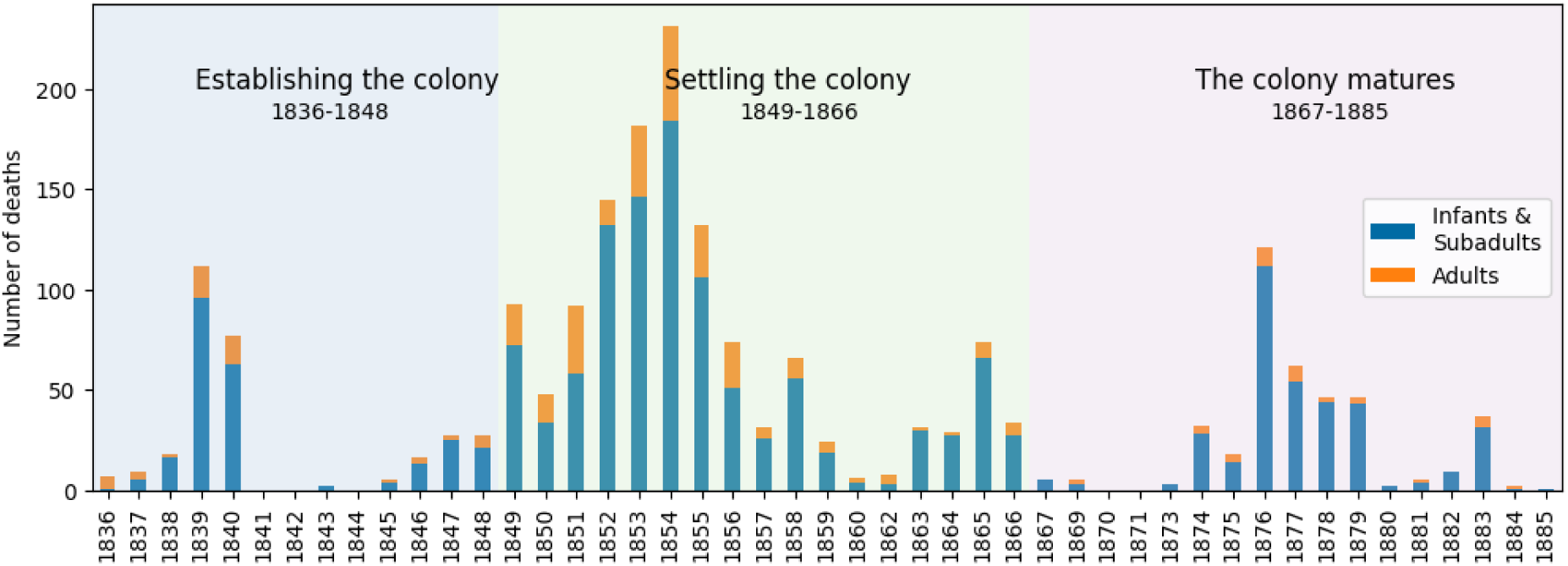
1836 - 1885. Number of infant and subadult, and adult deaths by year as recorded on emigrant voyages to South Australia from the United Kingdom, broken down by age group. [51, 52].

Table 4 provides the findings for the ships with high mortality rates on emigrant voyages from the UK to SA from 1836-1885 [51, 52]. It presents a breakdown of the number of passengers and deaths on each ship, along together with the percentage of fatalities recorded.

**Table 4.** 1836 -1885. Emigrant Ships by year of departure with a high percentage of deaths at sea carrying more than ten passengers from the United Kingdom to South Australia. (State Records of South Australia, 2016, 2022).

| Name of Ship | Year of Departure | Number of Passengers | Number of deaths | Death % |
| --- | --- | --- | --- | --- |
| Prince Regent | 1839 | 199 | 20 | 10% |
| Resource | 1839 | 211 | 30 | 14% |
| Fairfield | 1840 | 186 | 35 | 19% |
| Himalaya | 1849 | 176 | 29 | 16% |
| Douglas | 1850 | 125 | 21 | 17% |
| Shackamaxon | 1852 | 696 | 65 | 9% |
| John Bunyan | 1854 | 340 | 29 | 9% |
| James Fernie | 1854 | 376 | 30 | 8% |
| Morning Star | 1862 | 452 | 26 | 6% |
| Lochee | 1877 | 524 | 26 | 5% |

#### Superintendent Surgeons’ Logs - Causes of Death at Sea - 1849–1865

Investigation of the Superintendent Surgeons’ logs for the period 1849-1865 provides the number of deaths at sea for the study period. These data are divided into different age groups: 0-4 years, 5-18years and 19-50+ years. Infants and younger subadults were impacted the most during the voyage as highlighted in Fig. 5.

**Fig. 5.**
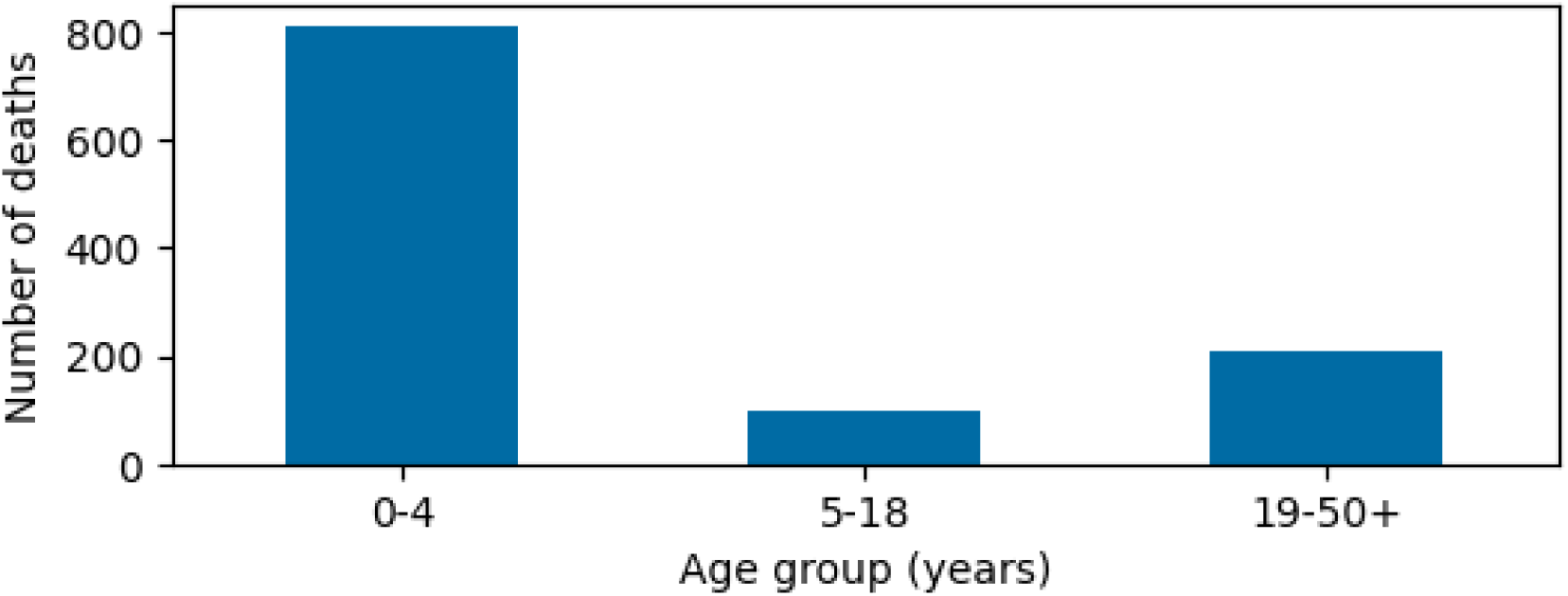
1849-1865. Number of deaths by age range in years for passengers on board emigrant ships to South Australia from the United Kingdom [54].

The death of a passenger at sea (1849-1865) (Government of South Australia, 1866), could have been caused by multiple interacting factors including infectious and non-infectious conditions. Table 5. lists the frequently recorded non-infectious and infectious conditions for all age groups on emigrant voyages from the UK to S A during this period.

**Table 5.** 1849 – 1865. Non-infectious and infectious conditions recorded for all age groups on emigrant voyages from the United Kingdom to South Australia [54].

| Common causes of death at sea |  |
| --- | --- |
| Non-infectious conditions/<br>Symptoms | Example/s of recorded causes |
| Fever | Allergic reaction, sunstroke |
| Cerebral | Hydrocephalus; effusion of the brain, meningitis |
| Chest | Heart disease/ failure |
| Abdomen - Liver, Kidney, Bowel | Organ failure, mesenteric disease – (small intestine) |
| Wasting & deficiency | Sea sickness, vomiting, atrophy |
| Convulsions | Epilepsy |
| Debility | Decay of strength |
| Marasmus | Progressive emaciation - malnutrition |
| Maternal | Miscarriage, haemorrhage |
| Neonatal | Premature birth, stillbirth |
| Croup | Coughing, inability to breath |
| Accident- trauma | Falling overboard, crushed |
| Suicide | Jumping overboard |
| Natural causes | Found dead |
| No cause recorded | N/A |
| Infectious conditions/<br>Symptoms | Example/s recorded causes |
| Fever | Viral, bacterial |
| Diarrhoea, Dysentery | Intestinal parasites, bacterial, viral |
| Putrid sore throat | Cynanche maligna; quinsy |
| Viral infection | Rubella, measles, smallpox |
| Bacterial infection | Scarlet fever, scarlatina, cholera, typhoid typhus, teething |
| Upper respiratory | Phthisis, tabes mesenterica, whooping cough, bronchitis, pneumonia |
| Maternal | Puerperal -postpartum infection |
| Sexually transmitted | Syphilis |

Documentation for 142 emigrant ships that sailed from the UK to SA (1849 to 1865) show that from the 89165 passengers recorded in voyage records during this period, 1141 deaths were logged. Therefore, around 1.3% or 1 of every 80 died before arrival [54]. From the total number of deaths, 387 (34%) have no cause or explanation recorded. Analysis of this data is further complicated due to a variety of spellings and terms used to record the cause of death [54].

The most commonly documented ‘causes’ of death for each age group on board of the 142 voyages are given in Table 6. These are shown in the order of the frequency that they were recorded in the ships’ logs e.g., numerous infants under one year of age died due to diarrhoea or dysentery followed by ‘debility’, convulsions and bronchitis. Fig. 6 breaks down this data further showing the number of deaths for each ‘cause’ and in which year/s had the highest number or a ‘peak’ in the recorded deaths on emigrant voyages to South Australia from the United Kingdom from 1849-1865.

**Fig. 6.**
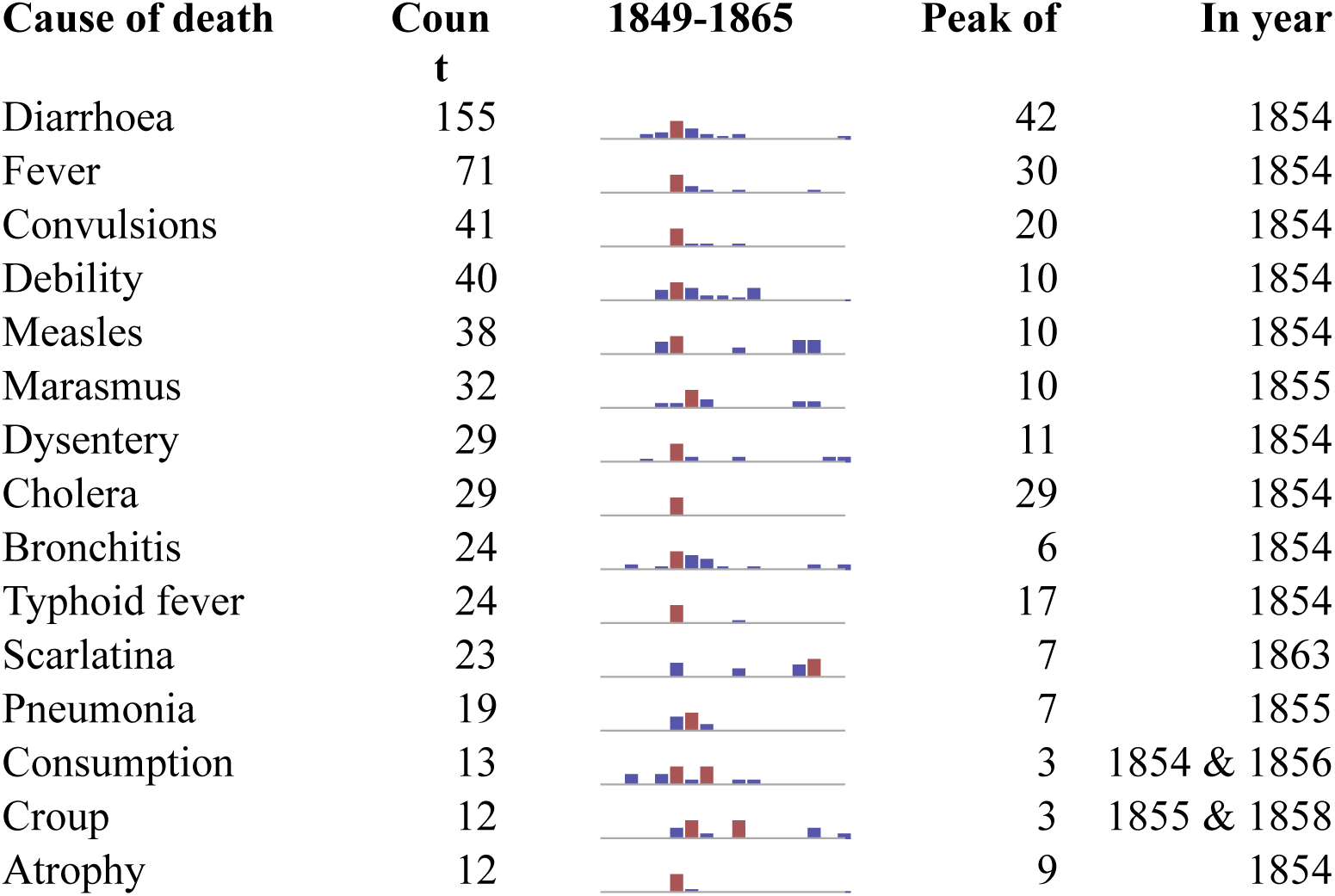
Top 15 causes of death on emigrant voyages to South Australia from the United Kingdom from 1849-1865 (Total Deaths: 1141) [54].

**Table 6.**
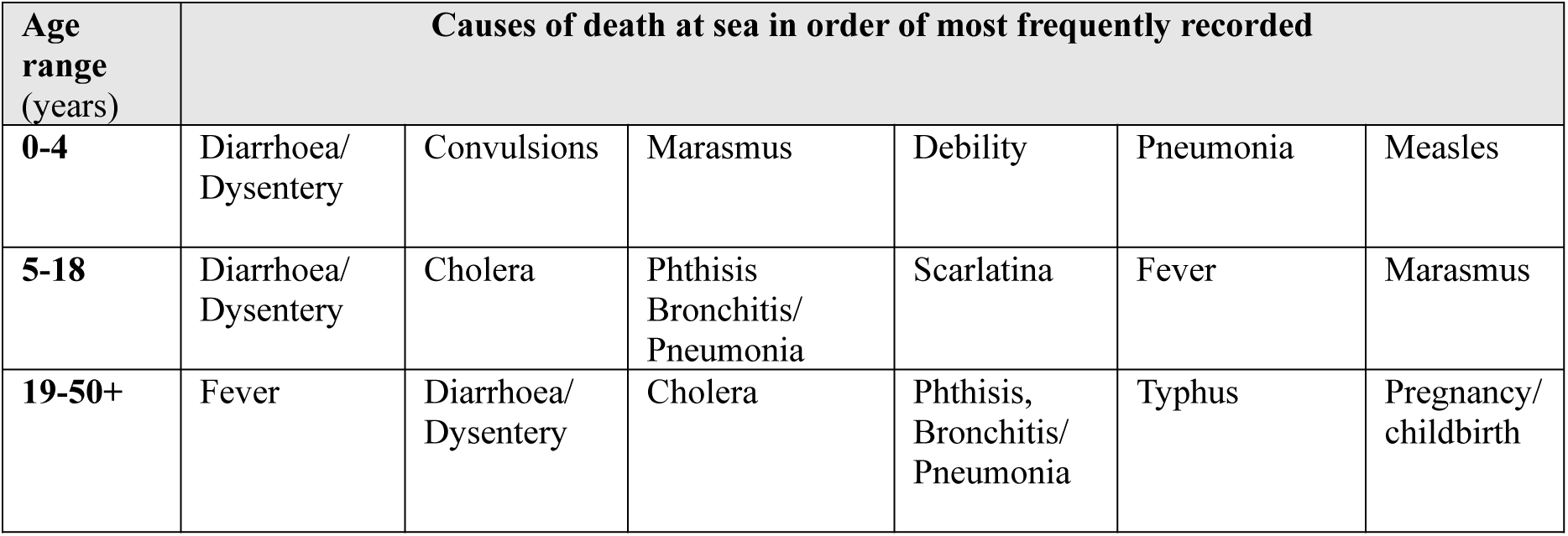
1849-1865. Causes of death at sea in order of most frequently recorded by age group on migrant ships from the United Kingdom to South Australia [54].

### 3. Church burial records

Burial records for St Mary’s Anglican Church, SA, show that a group of 143 individuals *could* have been interred in the unmarked ‘free ground’ area of the cemetery(see materials for details) [55]. These burials, paid for by the SA government, fluctuated between 1847 and 1885 (Fig. 7), with the majority of them taking place before the 1870s. The number of internments in the unmarked section declined after this time i.e., the 1870s (Fig. 7) [43, 55]. The highest number of deaths/burials for this group of 143 individuals where amongst the infants - age range 0-11 months (67/143), followed by the subadults – age range 1-3 years (38/143), (Table 7) [43].

**Fig. 7.**
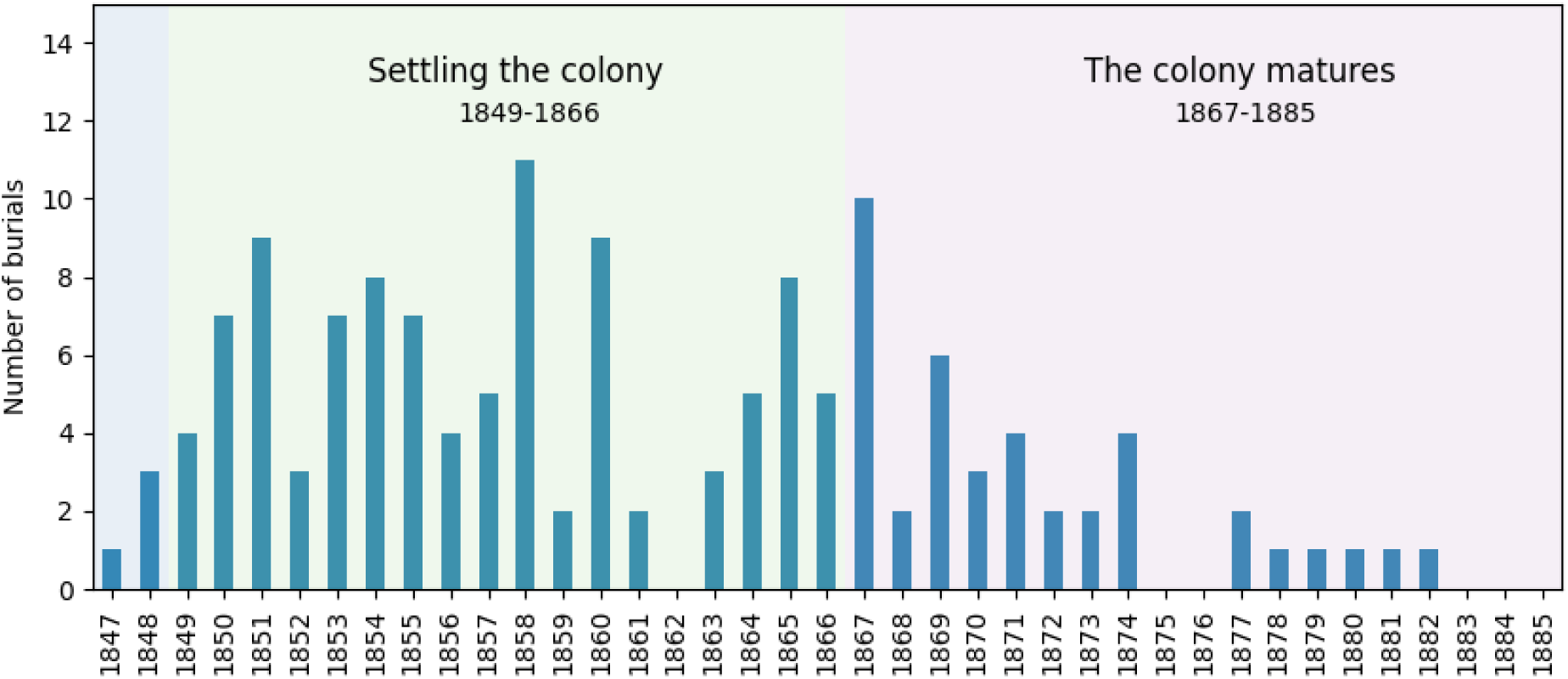
1847-1885. St Mary’s Church Burial Records -Year of burial and number of individuals – (total N=143) listed as interred in the unmarked section of St Mary’s Cemetery, South Australia, (or the location of their burial was not recorded, or is unknown due to damage to records) [55].

**Table 7.** 1847-1885. St Mary’s Cemetery Records– Most frequently recorded causes of deaths with age ranges of the deceased [43].

| Commonly Recorded Cause of death | Age range - years |  |  | Number of individuals listed |
| --- | --- | --- | --- | --- |
|  | Infant / young subadult 0-4 | Subadult 5-18 | Adult 19-50+ |  |
| Unknown Causes | 28 | 2 | 5 | 35 |
| Dysentery, Gastritis | 18 | 0 | 0 | 18 |
| Atrophy | 11 | 0 | 0 | 11 |
| Teething | 9 | 0 | 0 | 9 |
| Pulmonary conditions | 2 | 1 | 5 | 8 |
| Convulsions | 8 | 0 | 0 | 8 |
| Accident | 0 | 2 | 3 | 5 |
| Mesenteric disease | 4 | 0 | 0 | 4 |
| Disease of the brain, insanity | 1 | 0 | 2 | 3 |
| General decay/ old age | 0 | 0 | 3 | 3 |

Records frequently listed diarrhoea and dysentery as a cause of death for the St Mary’s infants and subadults (Table 7). Phthisis (tuberculosis) and natural decay were commonly listed as causes of death for some adult males, and accidents caused the death of three adult males from this group (Table 7). For the adult females, conditions related to the womb and the brain were listed (Table 7) [43].

## Discussion

### Integration of the multiple data sources

The integration of multiple data sources (Table 1) provided an insight into the morbidity and mortality rates for early migrants both at sea and in South Australia. Further, the multi-methodological approach used for the analyses of the dentition and other skeletal materials, provided an improved understanding of the health conditions experienced as well as an insight for some of the challenges faced by this specific group of migrants. In this way the application of the Wisdom Hierarchy and the CAS framework to the three diverse data sources provided new, more nuanced insights than would result from the independent treatment of each source.

The findings for the combined data sources showed a general trend for high infant and subadult mortality both at sea during the migrants’ voyage and after arrival and settlement during the early years of the colony. The young demographic profile of the individuals who represented the highest number of passenger deaths at sea (Fig. 4), was reflected in the burial records associated with St Mary’s Church’s (Table 7). This documentary trend was confirmed by the high number of infants and subadults excavated from the unmarked section of St Mary’s Church Cemetery (Table S1).

During the voyage to SA, for some passengers’, health issues such as sea sickness, and/or poor conditions onboard the ship, lack of ventilation, and contaminated food and water supplies would have increased their susceptibility to further health insults. The environment on board also meant that diseases such as measles and smallpox were likely to spread quickly due to the concentration of individuals in a confined space compared to on land.

In the colony during the early period (1836-1848), rudimentary health care was available in the city [58], but for the individuals living in communities away from these facilities reduced access to a doctor could prove fatal. The health of the younger age groups would have been impacted more than the adults due to their small body size and the reduction of the maternal antibodies combined with a still developing immune system. These factors would have reduced their ability to recover from the effects of disease such as gastrointestinal conditions and associated dehydration compared to an adult [59].

The background of the individuals in this study i.e., the excavated sample and those listed in St Mary’s burial records, is unknown, but the majority of them are likely to have been born in Britain. Some may have had a low socioeconomic status, working in a factory or as labourers. Having survived the voyage to South Australia, they went on to settle and experience a new lifestyle. Many individuals thrived in the colony, seizing new opportunities, building commercial enterprises or substantial landholdings. However, the financial position when they died of those buried in the unmarked ‘free ground’ section of St Mary’s Cemetery and the evidence from their skeletal remains show that they did not prosper in the early decades of this colony. Rather the challenges of the new life, the volatility of the economy and the limitations of the infrastructure proved overwhelming.

### Bioarchaeological investigation of St Mary’s Cemetery Sample

Poor oral health, such as extensive carious lesions, periodontal disease, evidence of pipe smoking and widespread antemortem tooth loss affected the majority of the adults in this skeletal sample (Table 3) [47]. This poor oral health would have negatively influenced the individual’s general systemic health. Dental developmental defects e.g., enamel hypoplasia and interglobular dentine (Table 3), seen in many subadults and adults confirmed that these individuals had suffered but survived one or more health insult during the development of their dentitions. Whether these conditions occurred before migration in the UK, during the voyage or in SA is unknown.

Evidence of the skeletal developmental defect sacral spina bifida occulta was observed in several individuals (Table 3). This condition is compatible with a lack of folic acid (Vitamin. B9), which leads to the failure of the vertebral arches of the sacrum to fuse and is linked to a deficiency in the pregnant woman’s diet [60]. This condition and other metabolic deficiencies e.g., lack of Vitamin C and/ or iron (Table 3), may have occurred due to limited access to necessary nutrients, education and/ or poor antenatal health care. These conditions highlight some of the many challenges that the migrants faced.

Morphological changes to the bones due to diseases such as vertebral osteophytes, Schmorl’s nodes, and eburnation of vertebral facets and/or other surfaces, were identified on several of the adults from this excavated sample (Table 3) [45, 46]. Bony outgrowths and/or changes to the structure of the joints would have caused pain and reduced the mobility of the sufferer and suggest that the affected individuals had undertaken hard physical labour over a long period of time. Changes to the bone/s due to infectious disease e.g., tuberculosis was also identified in the excavated skeletal remains (Table 3) [43, 56, 57]. All these factors, poor oral health, systemic diseases and deficiencies, and excessive demands of some types of hard physical labour, would have contributed to poor general health and shorter lifespans of this group.

### Ships Records

#### The Voyage – UK to SA

The number of voyages from the United Kingdom to South Australia between 1836 -1885 were influenced by multiple interacting factors including political instability, economic fluctuations and environmental events i.e., a dip in the number of ships arriving in SA were linked to the colony’s first economic depression, the multiple changes of colonial Governors and periods of drought (Fig. 3) [20, 38, 61, 62]. Increases in immigration trends were also closely tied to the colony’s evolving needs, for instance tradesmen, engineers, and builders were likely to be required to support infrastructure growth and development projects, whereas farm labours would have been needed to develop agricultural industries. Economic booms and crises prompted or reduced the number of emigrant voyages to the colony. British colonial policy, trade dynamics, and changes to the UK labour market could also influence the recruitment of migrants.

#### Superintendent Surgeon and documented causes of death at sea

Large emigrant ships travelling from the UK to SA could convey between 500-700 passengers at one time. This high number of people would mean there was limited space onboard, and that disease could quickly spread through a vessel [51, 52]. The cramped conditions, quality of nutrition and sanitation on the ship, as well as the competency of the Captain, Surgeon and the crew would have all impacted the health status of the passengers, with some of the early voyages having a high mortality rate (Table 5).

The system of appointing a Superintendent Surgeon to each emigrant ship carrying more than 100 passengers aimed to reduce mortality rates at sea [42]. Excessive deaths during the voyage meant not only financial penalties for the surgeon but also risked damaging his medical reputation and that of the Captain and British Government. High mortality rates could even deter future migrants from travelling to SA, with the roll-on effect of potentially reducing political, and economic gains.

Over time, the initial anticipated role of the Superintendent Surgeon evolved with emergent duties that encompassed numerous concerns and was not limited to attending sick and injured passengers. Dr George Mayo on board the emigrant ship ‘Asia’ [63, 64] was called upon to settle a dispute between passengers. He notes, “this morning, I was called to Mr Letts whom Mr. C. Olliver had struck and made his face bleed and as the complaint came it was done without provocation. I had the parties into the cabin before myself and the captain…” [64]. He also recorded concerns about the migrants’ attitude, their food provisions and health in his diary, “I have great trouble to get the emigrants on deck” and “The rotten potatoes in the hold I think occasioned dysentery amongst the emigrants, spoken to Captain. Fredeman.” [64]. These multiple duties are illustrative of non-predictive outcomes, both positive and negative and are consistent with a CAS framework.

The pattern of passenger deaths at sea by year (Fig. 4) did not closely follow that of the number of voyages per year (Fig.3) e.g., the years with the highest number of voyages (1849 and 1850) did not have the highest number of deaths. This suggests that other factors influenced the death rate at sea. On occasion, particular conditions such as diarrhoea or cholera could affect a large number of the migrants during a voyage (Fig. 6) [51, 52, 54]. It is possible that these deaths were linked to similar epidemics on land such as the Broad Street cholera outbreak in London during the same year [54, 65, 66]. Could contaminated water have been brought aboard this ship? While the limits of this data do not give a clear answer, many deaths recorded as caused by diarrhoea or cholera did occur in 1854 (Fig. 6) [54, 63]. There may have been concern that ships affected by such conditions arriving in SA would affect the population of the new colony. However, available SA newspaper accounts relating to voyages with recorded cases of cholera suggest no active cases disembarked and that cases with this condition had previously been diagnosed in the colony [67–70].

The age of an individual and their health status at the time of the voyage to SA meant that any infectious or non-infections conditions encountered on the ships could develop into fatal illnesses. Infectious and non-infectious conditions such as measles, whooping cough, and/ or consuming contaminated water, lack of cleanliness, and at times insufficient medical supplies on board have been recorded (Table 5) [51, 52, 54]. Other risks to the health and safety of the passengers during the voyage were connected to inadequate ventilation below decks, and flooding of their quarters. Severe weather conditions, leading to a shipwreck, were also a possibility [71].

Diarrhoea was recorded as a major cause of death for infants and young subadults by Superintendent Surgeons between 1849 and 1865 (Table 6) [54]. These infants and subadults under the age of 4 years represented the largest demographic on the voyage and the most vulnerable age groups due to their developing immune system and the reduction of the maternal antibodies that had protected them during their first year of life (Fig 4).

For many adults the major causes of death at sea were fever and ‘phthisis’ a term used in the 19^th^ century for tuberculosis [72] (Table 5). It is highly likely that some migrants were suffering from medical conditions before embarking, from which they subsequently died at sea [54]. The recorded cause of death in the ship’s log could have occurred due to multiple interacting conditions. The diagnoses recorded as diarrhoea and/or fever are symptoms rather than diseases; and may be indications of more than one underlying disease.

The loss of young individuals had a sub-generational impact on the forming colony in South Australia. In addition, the specific disease outbreaks at sea that were not fatal for the majority of subadults and adults may well still have had lasting health impacts.

The number of deaths on emigrant ships decreased over time suggesting an improvement in cleanliness, the storage of food and water, and a growing awareness of how diseases were transmitted (Fig. 4. and Table 4) [41, 73]. The reduction in mortality rates could also be related to amendments to the Passenger Act of 1835, as well as technological advances such as ventilation machines for the lower decks of the ships and distillation of water [65, 73–75].

#### Life in the new Colony – St Mary’s Burial Records

While some migrants overcame the challenges and flourished in the new Province of South Australia there were others who had difficulties [33, 76]. Complex dynamic interests that interacted to influence the colony’s development could have also influenced and affected the health of the population [20, 38, 62].

The arrival of emigrant ships with large numbers of passengers should have provided the colony with a varied work force, but the time delay involved in communicating between Adelaide and London often led to the arrival of substantial numbers of immigrants all with similar skills for whom there was no employment [77]. This could be due to the rapidly changing economy and added further to health and socioeconomic pressures. For example, records show that several groups of unmarried Irish women and girls arrived in SA, between 1848 -1856, perhaps with a promise of work ‘in service’ as a maid or housekeeper and a future far from famine [53, 78–80]. Their arrival, over a short period of time would have flooded the local labour market for domestic servants [80].

This unsustainable situation created by increased unemployment in the colony corresponds with a rise in the number of people seeking financial support from the Government and charitable organisations [81–86]. Official government returns for 1853 states that 464 people received relief from the Destitute Asylum, while in 1854 it was 685 people but by 1855 the number was 3,027 [61, 87]. As the colony matured a ‘Constitution for South Australia’ was developed (Fig. 3) and elections held for a new Parliament [88–90]. This enabled some of the populace to vote and the colony to exercise self-rule. South Australia was now starting to generate its own future workforce but during this period a change in the ‘Regulations for the selection of Persons in Britain for Free Passage’ to the colony in 1858 also reduced the number of emigrants arriving from the UK (Fig. 3) [90, 91]. These changes are seen in the returns for the SA census of 1860, which confirms that one third of the population had now been born in the colony [92]. The newly formed Parliament of South Australia was now exploring the issue of social security through the Destitute Person’s Relief Act of 1866/ 67 [93], and the advance of tertiary education with the foundation of the University of Adelaide in 1874.

#### Limitations of this Study

The data used and the findings of this study relating to migrant deaths at seas could be skewed as many of the emigrant ships from the UK to SA during the study period (1836- 1885) had no fatalities. However, the length of the journey, condition on route such as the weather and/or overcrowding onboard ship meant that there was loss of life on many voyages.

Limits of historical records associated with emigrant ships to SA, include variability in the details recorded in the Superintendent Surgeons’ logs. Analysis relating to the causes of death at sea was limited to 1849 to 1865 due to the nature of the sources themselves. Historical sources are rarely a complete and comprehensive record and may contain bias due to both individual perspective and prejudices of the broader political and social culture.

Damage to and loss of records over time was also a problem. A fire at St Mary’s Anglican Church destroyed many historical records stored in the building, which could have affected the availability of data relating to burials in the unmarked section of the church cemetery [94, 95]. There were limits regarding the extent to which information relating to the health of the excavated individuals could be collected from the skeletal remains. For example, no data relating to conditions that affect the soft tissues of the body could be gained.

## Conclusion

The Complex Adaptive Systems approach increased knowledge of the general health of 19^th^ century migrants from the UK to SA and has resulted in additional insights that could not have been gained using a single analytical methodology.

This methodology provided a deeper understanding of the emergent and unpredicted outcomes that occur with the interaction of multiple factors in varied data sources related to a rare colonial South Australian skeletal sample and diverse historical texts. This study demonstrates that a multidisciplinary/ multi-methodological/ CAS approach and analysis can now be undertaken to explore the identities and life histories of specific individuals in this sample. It can be further developed and applied to other archaeological and modern samples.

## Acknowledgements

The authors wish to thank: Dr Kiera Lindsey, South Australia’s History Advocate, for her encouragement of this study and her expert advice related to the history of this early colony. Rev. Canon William Deng, Priest of St Mary’s Anglican Church, South Australia, who provided access to the St Mary’s skeletal sample and the parish records. As indicated in the text of this paper the excavation of the skeletal sample was undertaken by Dr Timothy Anson in 2000 under the supervision of Prof. Maciej Henneberg and Emeritus Prof. Don Pate.

The authors acknowledge the instruments and expertise of Microscopy Australia (ROR: 042mm0k03) at Adelaide Microscopy, University of Adelaide, enabled by NCRIS, university, and state government support. Dr Agatha Lambrinidis provided assistance with the Bruker SkyScan 1276 Small Volume Micro-CT system. Associate Prof. Egon Perillli and Dr Sophie Rapagna from the Medical Device Research Institute, College of Science and Engineering, Flinders University, shared their expertise regarding the Large Volume Micro-CT system. Flinders Microscopy and Microanalysis (FMMA) provided access to the Nikon XT H 225 Micro-CT scanning system. The Australian Research Council (LE180100136) provided funding contribution for the procurement of the Large Volume Micro-CT system.

## Supporting Information

**Table S1.** 1847-1927 – Demographic profile: Estimation of age range from the analysis of dentition and skeletal remains with the number of individuals within the different age ranges. Estimation of sex with the number of individuals.

